# Systematic exploration of protein conformational space using a Distance Geometry approach

**DOI:** 10.1101/650903

**Authors:** Thérèse E. Malliavin, Antonio Mucherino, Carlile Lavor, Leo Liberti

**Author notes:** Corresponding authors Thérèse E Malliavin, Institut Pasteur and CNRS UMR 3528, Unité de Bioinformatique Structurale, 25 rue du Dr Roux, 75015 Paris, France E-mail address, Leo Liberti, CNRS LIX (UMR 7161), Ecole Polytechnique, 91128 Palaiseau, France.

## Abstract

The optimisation approaches classically used during the determination of protein structure encounter various diffculties, specially when the size of the conformational space is large. Indeed, in such case, algorithmic convergence criteria are more difficult to set up. Moreover, the size of the search space makes it difficult to achieve a complete exploration. The interval Branch-and-Prune (iBP) approach, based on the reformulating of the Distance Geometry Problem (DGP) provides a theoretical frame for the generation of protein conformations, by systematically sampling the conformational space. When an appropriate subset of inter-atomic distances is known exactly, this worst-case exponential-time algorithm is provably complete and fixed-parameter tractable. These guarantees, however, immediately disappear as distance measurement errors are introduced. Here we propose an improvement of this approach: the threading-augmented interval Branch-and-Prune (TAiBP), where the combinatorial explosion of the original iBP approach arising from its exponential complexity is alleviated by partitioning the input instances into consecutive peptide fragments and by using Self-Organizing Maps (SOMs) to obtain clusters of similar solutions. A validation of the TAiBP approach is presented here on a set of proteins of various sizes and structures. The calculation inputs are: a uniform covalent geometry extracted from force field covalent terms, the backbone dihedral angles with error intervals, and a few long-range distances. For most of the proteins smaller than 50 residues and interval widths of 20°, the TAiBP approach yielded solutions with RMSD values smaller than 3 Å with respect to the initial protein conformation. The efficiency of TAiBP approach for proteins larger than 50 residues will require the use of non-uniform covalent geometry, and may have benefits from the recent development of residue-specific force-fields.

## Introduction

Since the early days of structural biology, optimization techniques have been at the heart of biomolecular structure calculation. Indeed, most of the experimental information is only indirectly related to protein structure. In addition, this information is noisy. Furthermore, the sparsity of data is made even bigger as most of biophysical techniques concentrates on time-average or space-average data in order to obtain large enough signal-to-noise ratio.

Several optimization schemes have been used for Nuclear Magnetic Resonance (NMR) structure determination, such as simulated annealing^1^ and genetic algorithms.^2^ Nowadays, several approaches exist for protein structure determinations by NMR.^3–7^ A Bayesian approach,^8^ using a Markov chain Monte Carlo (MCMC) scheme for the conformational space sampling,^9,10^ allowed the increase of the convergence radius for problems of protein structure determination by NMR. Furthermore, the use of a log-harmonic shape for distance restraint potential,^11^ along with a Bayesian approach for the restraint weighting,^12^ allowed an improvement of the quality of NMR protein structures.^13–15^ Log-harmonic restraints defined using Bayesian inference have been also recently^16^ proposed for back-mapping from coarse-grained models to atomic structures.

Most of the optimization methods used so far provide no guarantees of optimality, although they are commonly used in the hope of obtaining the global minimum or several global minima of the optimization problem. This, however, depends on the choice of a starting point for the computation. Consequently, calculations of protein conformations under NMR restraints are repeated several times during the procedure of structure determination,^17^ and the convergence of these calculations is generally required in order to accept a set of conformations as a solution. This iterative frame, ^6,18^ however, encounters difficulties when the problem has many local minima that are far apart. Such cases started to occur more frequently in the field of structural biology with the growing interest to disordered regions of biomolecules.^19–21^ Monte Carlo approaches have been proposed for intrinsically disordered proteins^22–24^ and molecular dynamics simulations^25^ are also used on all kind of biomolecular polymers, but they do not provide a definitive answer to the problem of finding all minima.

Since NMR studies biomolecules in solution, and due to the large number of various parameters it can measure, it is particularly sensitive to the effect of internal mobility. NMR measures are inter-atomic distances and angles, which are closely related parameters. The problem of protein structure determination by NMR can be thus considered as a Distance Geometry Problem (DGP).^26,27^ The interval Branch-and-Prune (iBP) approach has been developed^28^ for solving the DGP in the framework of the calculation of protein conformations. In this approach, an atom re-ordering^29,30^ ensures that there is a restricted and manageable locus for the spatial position of every atom. This is achieved by using a “relaxed form” of trilateration with respect to the three preceding atoms in the order. More precisely, two out of three of the distances involved in trilateration must be known exactly, and one may be subject to uncertainty and represented by an interval. Any atom, together with its three reference predecessors, give rise to a 4-clique in the protein graph: the iBP approach mimics the approach of exploring protein conformation in torsion angle space. ^31–34^ In the clique, exact distances are provided by covalent bond lengths and bond angle values, using the cosine law. Note that applying this framework using generic information from a force field instead of measured distances makes the implicit assumption of a uniform covalent geometry within the protein structure. Analyses of high-resolution crystallographic structures, ^35,36^ however, have shown that this assumption is not necessarily verified. Independent parallel work has conducted to the development of residue-specific force field.^37–40^

Basing on the atom reordering, it is possible to describe a tree exploration algorithm in order to find all solutions of a DGP instance. Each tree node represents a spatial position for an atom. The level of a node in the tree is the index of the atom in the reordering. This implies that all positions on the same tree level are the possible spatial positions for the atom having as rank this level. The width of the tree increases exponentially in the worst case, but it can be bounded to more manageable levels^41^ by choosing specific atomic orders. This yields a fixed-parameter tractable behavior (at least with exact distances). We note that the exploration of this tree is complete but implicit, in the sense that certain sub-trees are pruned because the atomic positions at their root nodes are not consistent with long range distances to preceding atoms. Naturally, each pruned node induces the pruning of the sub-tree rooted at that node. It was demonstrated^27,28^ that, starting from a set of exact distances measured in a given PDB structure, the search tree can be completely explored in a relatively small amount of CPU time.

Here, we employ the iBP algorithm in a setting which is considerably closer to the protocols of protein structure determination than the mathematical setting in which it was initially conceived. Instead of exact distances measured on a given PDB structure, this requires the use of a mixed set of distance intervals and of exact distances arising from a covalent geometry defined through a force field. Several attempts have been made in this direction in the recent past. A significant exploration of the conformational space of some *α*-helical 15 to 51-residues proteins was performed in Ref.,^42^ and more recently, the iBP approach was re-implemented^43^ in order to allow its application to real-life cases of protein structure determination. First, the number of tree branches was reduced^44^ by taking into account the information from improper angles. Second, a parser and a grammar have been defined to convert the topology, parameter and atom type information used in molecular modeling to the distance information which is the main input of iBP. Third, a syntax has been defined to make the atom reordering information a user-defined input of the calculation. This new implementation makes it possible to perform tree branching on intervals determined on *ϕ* and *ψ* backbone angles, which may be obtained through chemical shift measurements.^45^ Nevertheless, no systematic exploration of the protein conformational space has been previously attempted.^43^

In the present work, we employ the implementation of Ref.^43^ to develop a new strategy which allows a systematic sampling of the conformational space of small proteins and we validate this strategy using a set of various protein structures. The expected combinatorial explosion is prevented by several ingredients: (i) the division of the protein into fragments which are sampled independently and then assembled, (ii) the extensive use of signed improper angle values to reduce the tree size of each fragment, (iii) the use of self-organizing maps to cluster conformations of intermediate fragments.

The geometrical information used for input calculation corresponds to relevant NMR measurements on proteins. Indeed, NMR chemical shifts are easily measurable parameters. The relationship between chemical shifts and atomic coordinates is not straightforward, but several methods, as the neural network TALOS-N^45^ or chemical shift prediction approaches,^46–48^ exist for relating chemical shifts and atomic coordinates. Among them, TALOS-N ^45^ predicts from chemical shift values, (*ϕ, ψ*) likelihood distributions. The existence of such distributions supports the use of intervals on *ϕ* and *ψ* values as inputs for the TAiBP approach. In addition to the *ϕ* and *ψ* intervals, distance restraints with interval widths of 6 and 10 Å and defining qualitatively the protein global shape were used as inputs.

The proposed approach is called threading-augmented interval Branch-and-Prune (TAiBP) approach, as it intends to generate conformations of peptide fragments using iBP, as well as to thread these fragments in 3D space in order to build protein conformations. The name was coined in analogy to the threading approach^49^ used in protein 3D structure prediction. We point out that the idea to separate iBP instances in sub-instances is not completely new, but it was explored so far only in the context of parallel^50^ and distributed^51^ computing. Also, the idea of constructing protein conformations from fragment assembly was proposed initially^52–54^ in the Rosetta approach for protein structure modeling.

The proposed methodology is innovative with respect to the state of the art because it is designed to find all possible configurations compatible with a given set of angle and distance restraints on a given protein. This is in contrast to classical methods for structure determination,^1^ which might at best produce different protein conformations. The approach is different with respect to the more recently proposed methods aiming at determining the global minimum configuration of the system^4,55–59^ or at determining all relative positions of monomers within a protein homo-oligomer.^60^ On the contrary, the exhaustive list of conformations generated by TAiBP provides solutions for a larger range of problems.

It is important to note that our purpose is beyond finding a conformation close to the target one, since we aim instead to the much more ambitious goal of finding many (and hopefully all) incongruent but geometrical consistent conformations. Moreover, because our algorithm approach is not iterative but based on branching, we have no need for considering “convergence to a local optimum” a requirement for accepting a conformation. The results of our computational experiments, however, have been validated by detecting whether conformations close to the target PDB structure have been sampled during the tree exploration, by Root Mean Square Distance (RMSD) of atomic coordinates to the target structure. The proposed approach allows us to explore the tree for proteins up to 50 residues. The non-uniform covalent geometry, prevents (by now) our method from being successful on proteins larger than 50 residues.

## Materials and Methods

### Test case database

The database of protein structures was built in the following way. The protein structures contained in kinemage.biochem.duke.edu/databases/top100.php^61^ have been downloaded. This database was chosen as high resolution X-ray crystallographic structures on which hydrogens have been added with rotational optimization of OH, SH and NH^3+^ positions,^61^ thus producing objects corresponding to those iBP is designed to calculate.

Protein structures with number of residues between 21 and 107 have been selected, containing only trans peptidic bonds and corresponding to the following list of 24 proteins: 1aacH, 1benABH, 1bkfH, 1bpiH, 1ckaH, 1cnrH, 1ctjH, 1difH, 1edmBH, 1fxdH, 1igdH, 1iroH, 1isuAH, 1mctIH, 1ptfH, 1ptxH, 1rroH, 256bAH, 2bopAH, 3b5c, 3ebxH, 451cH, bio1rpoH and bio2wrpH. In 3b5c, the N terminal residue T88 was removed because of missing backbone atoms. On each structure, the conformation of chain A was selected for preparing the iBP input, and in the case multiple conformations have been observed for a residue, the A conformation was selected.

### Input values for the calculation

The parameters defining the covalent and improper geometries were taken from the geometric force field PARALLHDG (version 5.3) ^62^ (Table 1). One should notice that, although these parameters were proposed more than two decades ago, they still correspond to the state-of-art of molecular force fields with fixed charges, as the covalent bond lengths and bond angles of most of fixed-point force fields were determined at the same time or earlier.^63^ The atom re-ordering is the same proposed than in the most recent implementation of iBP^43^ (Table 2).

**Table 1:** Geometric parameters for covalent and improper bonds and angles taken from the force field PARALLHDG (version 5.3). ^62^

| Atom name | Definition of atom type | Bond atoms | Bond length (Å) |
| --- | --- | --- | --- |
| CH1E | $\alpha$ carbon | C-CH1E | 1.525 |
| C | carbonyl carbon | CH1E-HA | 1.080 |
| O | carbonyl oxygen | CH1E-NH1 | 1.458 |
| OC | C-terminal carbonyl oxygen | CH1E-NH2 | 1.486 |
| HA | Hα hydrogen | C-NH1 | 1.329 |
| NH1 | amide nitrogen | C-O | 1.231 |
| NH2 | N-terminal amide | C-OC | 1.249 |
|  | nitrogen | H-NH1 | 0.980 |
| H | amide hydrogen | H-NH2 | 0.980 |
| Bond angle atoms | Bond angle value (°) | Improper angle atoms | Improper angle value (°) |
| C-CH1E-HA | 108.9914 | C-CH1E-HA-HA | -70.4072 |
| C-CH1E-NH1 | 111.1396 | CH1E-C-NH1-HA | 66.2535 |
| C-CH1E-NH2 | 106.9610 | C-NH1-HA-HA | -70.8745 |
| C-NH1-CH1E | 121.6541 | HA-CH1E-HA-HA | -66.5692 |
| CH1E-C-NH1 | 116.1998 | CH1E-C-NH1-CH1E | 178.0 |
| CH1E-C-O | 120.8258 | O-C-NH1-H | 178.0 |
| CH1E-NH1-H | 119.2367 | C-CH1E-NH1-O | 0.0 |
| C-NH1-H | 119.2489 | NH1-CH1E-C-HA | 119.0 |
| C-NH2-H | 118.1853 | NH2-CH1E-C-HA | 119.0 |
| HA-CH1E-NH1 | 108.0508 | NH1-HA-CH1E-C | 121.0 |
| H-NH2-CH1E | 109.5000 | NH2-HA-CH1E-C | 116.0 |
| H-NH2-H | 107.3000 | C-NH1-CH1E-H | 180.0 |
| NH1-C-O | 122.9907 | CH1E-C-NH1-O | 180.0 |
| NH2-CH1E-HA | 108.4800 | NH2-H-H-CH1E | 41.0 |
| NH2-C-O | 122.6277 | CH1E-OC-C-OC | 178.0 |
| CH1E-C-OC | 118.0611 |  |  |
| OC-C-OC | 123.3548 |  |  |

**Table 2:** Atom re-ordering used during the iBP calculation step within the first, the last and the inner residues of the peptide fragment. The order is described by the list of atoms names, the signs “-” and “+” describing atoms located in the previous and the next residues in the primary sequence.

| Residue position | order |
| --- | --- |
| first | N, H1, H2, CA, N, HA, CA, C |
| inner | N, -O, -CA, -C, N, CA, C, +N,<br>-C, N, CA, H1, N, CA, C, HA, C, CA |
| last | N, -O, -CA, -C, N, CA, C,<br>-C, N, CA, H1, N, CA, C, HA,<br>C, CA, O, C, O2 |

Two sets of values were used for the backbone angles *ϕ* and *ψ* in order to evaluate the effect of assuming a uniform covalent geometry on the TAiBP results.

i. the *ϕ*_angl_, *ψ*_angl_ angles of residue *i* measured on the X-ray crystallographic structures as the angles between planes C^*i-*1^N^*i*^C*α*^*i*^ and N^*i*^C*α*^*i*^C^*i*^ and between planes N^*i*^C*α*^*i*^C^*i*^ and C*α*^*i*^C^*i*^N^*i*+1^ using VMD.^64^ The plane ABC is defined as the plane passing through the positions of the atoms A, B and C.
ii. the *ϕ*_dist_, *ψ*_dist_ angles calculated from the distances d(N^*i*^,N^*i*+1^) and d(C^*i*^,C^*i*+1^) between N and C atoms of successive residues, assuming the covalent geometry uniform and described in Table 1. The dihedral or pseudo-dihedral angle Ω between ordered atoms *i*-3,*i*-2,*i*-1 and *i*, is determined using the cosine law from a trihedron:^44^

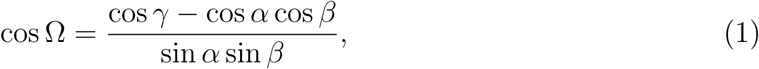

where *α* is the angle between atoms (*i*-3,*i*-2,*i*-1), *β* is the angle between atoms (*i*-1,*i*-2,*i*), and *γ* is the angle between atoms (*i*-3,*i*-2,*i*). For Ω angles being *ϕ* or *ψ*, the angles *α, β* and *γ* are calculated from the bond lengths and bond angles among heavy backbone atoms, as well as the distances d(N^*i*^,N^*i*+1^) and d(C^*i*^,C^*i*+1^) between successive residues along the protein sequence. The calculation is described in details in the Supporting Information.

The input restraints for iBP processing of peptide fragments are: (a) the restraints corresponding to the bond lengths and bond angles of the force field PARALLHDG (version 5.3);^62^ (b) the backbone angles *ϕ* and *ψ*, determined as described previously; (c) the distances between C*α* atoms located at the two extremities residues of each peptide fragment. The input restraints for the fragment assembly are: (i) the long-range distances between C*α* atoms of the residues located at the middle of each fragment; (ii) pruning devices avoiding that C*α* atoms belonging to different fragments are closer than 1 Å. The following error bounds have been used: errors of ± 10°, 20° and 30° for the angles *ϕ* and *ψ*, an error of ± 3 Å for the C*α*-C*α* distance between extremities of peptide fragments, and error of ± 5 Å for the long-range C*α*-C*α* distance between peptides fragments. Examples of inputs in tbl format are given in the Supporting Information.

### Interval Branch-and-Prune calculation of peptide fragments

As the TAiBP approach intends to explore the conformation of protein backbone, the processed protein is initially converted to a poly-Alanine chain. The protein is then divided in 15-residues peptide fragments, two successive fragments having a sequence of 5 superimposed residues. This fragment size was determined as it permits to obtain tree sizes which are manageable to explore in the reasonable amount of time, as it will be shown below in the subsection “Exploring the conformational space of fragments using iBP” of Results. The number of superimposed residues was chosen to avoid artifacts due to superimposition. The peptide fragments are assembled together to produce protein conformations, as it will be described in the next subsection.

For each fragment, the iBP tree of possible conformations is systematically explored. We employ the most recent implementation of iBP^43^ (in the C programming language), which is tuned for the calculation of protein conformations based on the force field knowledge for the covalent geometry. The tree branching is performed on the *ϕ* and *ψ* backbone angles. No branching was performed on the peptidic angle *ω*. Indeed, analyses of variations of *ω* angles in the X-ray crystallographic structures^65–67^ show that the angles *ω* mostly vary in intervals of ±8° around −180 and 180° which are smaller than the intervals sampled in the present work for *ϕ* and *ψ*.

The position of the atoms were determined as described in Ref.^44^ For any atom *i* in the order, we seek the atomic coordinates **x**_*i*_, given distances between the three preceding atoms *i -* 1, *i -* 2 and *i -* 3. As described above, the distances *d*_*i,i-*1_, *d*_*i,i-*2_ and *d*_*i,i-*3_ between atoms *i,i -* 1, atoms *i,i -* 2 and atoms *i,i -* 3 are known, where distance *d*_*i,i-*3_ between atoms *i* and *i -* 3 is potentially an interval. The variables *d*_*i*_, *θ*_*i*_ and *τ*_*i*_, where *d*_*i*_ denotes *d*_*i,i-*1_, permes to determine the position of atom *i* by the following equation:

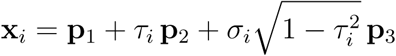

where **p**_1_, **p**_2_, **p**_3_ ∈ ℝ3 depend only on **x**_*i-*1_, **x**_*i-*2_, **x**_*i-*3_, *d*_*i*_ and *θ*_*i*_,

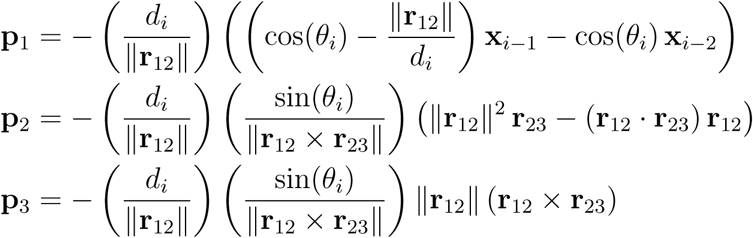

and we have introduced the scalars **r**_12_, **r**_23_ for notational simplicity,

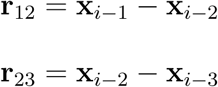

The angle *θ*_*i*_ is obtained from the cosine law using the relevant distances,

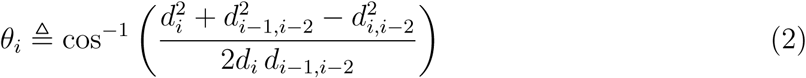

The pseudo-dihedral angle *ω*_*i*_ formed by the atoms *i -* 3,*i -* 2,*i -* 1 and *i* is partially determined by its cosine value cos *ω*_*i*_ = *τ*_*i*_, which is calculated using the cosine law for a trihedron:^44^

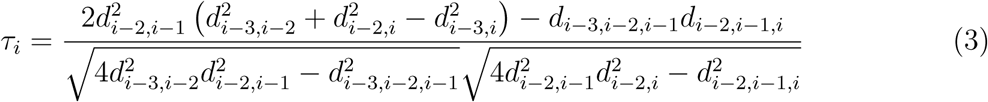

where

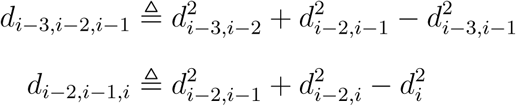

The determination of the pseudo-dihedral *ω*_*i*_ is completed by the sign *σ*_*i*_ ∈ {*-*1, +1} of sin *ω*_*i*_. When *ω*_*i*_ is known from either protein chemistry or measurement, we may directly compute *τ*_*i*_, as well as the sign *σ*_*i*_ ∈ {*-*1, +1}. This is the case when the angle *ω*_*i*_ corresponds to an improper angle (Table 1) and this allows to reduce the branching to one branch.

The number of saved conformations is reduced by applying a RMSD filter of 3 Å between two successively saved conformations. In order to avoid pruning due to slight discrepancy between distance restraints, a tolerance of 0.05 Å has been added to the bounds of distance intervals. The minimum discretization factor, which is the minimum ratio between each distance interval to the number of tree branches generated within the interval, was set to 0.05 Å, in order that the branching does not over-sample small intervals. No pruning due to the van der Waals radii of the force field protein-allhdg5-4 PARALLHDG (version 5.3)^62^ was applied. A maximum number of saved conformations of 10^9^ was permitted for each iBP run. The solutions are stored in a multiframe dcd format. ^68^

### Assembling the peptide fragments and clustering

The generated conformations of neighbouring peptide fragments in the protein sequence are then assembled by superimposing the five last and initial residues of the fragments located first and second in the sequence. The conformations of fragments are assembled by root-mean-square superimposition of backbone atoms located in the five superimposed residues. For each superimposition, the residue number for which the smallest distance was observed between corresponding atoms in the two peptides is used to decide where to stop with the first peptide and to continue with the second one. The assembled conformation is then submitted to two pruning devices: (i) a device checking whether there is no clash between the two fragments, i.e. no C*α* atoms closer than 1 Å, (ii) a device checking that long-range C*α*-C*α* distance restraints between peptide middle residues are verified. The fragment assembly is implemented using python scripting based on the MDAnalysis^69,70^ and numpy^71^ python packages.

To scale down the combinatorial explosion of the calculation, a clustering approach, the **S**elf-**O**rganizing **M**aps (SOM),^72–75^ which is an artificial neural network (ANN) trained using unsupervised learning, were used to reduce the number of conformations.^76^ The SOM displays the advantage with respect to the k-means clustering approach that it does not require the predetermined knowledge of the number of clusters. The SOM approach was used after a iBP calculation or after an assembly step as soon as the number of saved conformations was larger than 1000. The conformations sampled by iBP were encoded from the distances *d*_*ij*_ calculated between the *n* C_*α*_ atoms of the fragment, by diagonalizing the covariance matrix *C*:

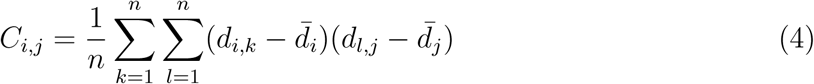

where 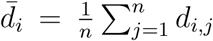. The information contained in the matrix *C* is equivalent to its four largest eigenvalues along with the corresponding eigenvectors. The eigenvalue and eigenvector descriptors are used to train a periodic Euclidean 2D self-organizing map (SOM), defined by a three-dimensional matrix. The first two dimensions were chosen to be 100 × 100 and define the map size.

The self-organizing maps were initialized with a random uniform distribution covering the range of values of the input vectors. At each step, an input vector is presented to the map, and the neuron closest to this input is updated. The maps are trained in two phases. During the first phase, the input vectors are presented to the SOM in random order to avoid mapping bias with a learning parameter of 0.5, and a radius parameter of 36.^77^ During the second phase, the learning and radius constants are decreased exponentially from starting values 0.5 and 36, respectively, during 10 cycles of presentation of all the data in random order. Once the calculation of the SOM has been realized, the conformations corresponding to local maxima of homogeneity, are detected and the total set of conformation is replaced by these representative conformations.

## Results

### Probing the hypothesis of uniform covalent geometry

The 24 structures extracted from the database of Word et al^61^ have been processed to analyze the geometry of covalent angles (Figure 1a). The distributions of covalent angles between C-N-C*α* (blue curve), N-C*α*-C (magenta curve) and C*α*-C-N (green curve) (Figure 1a) are centered on 121.3°, 110.6° and 116.8°, with standard deviations of 2.2°, 3.0° and 2.2°. These distributions agrees with the ones observed by Hinsen et al:^78^ C-N-C*α* (121.4^°^ ± 1.6°), N-C*α*-C (111.1° ± 2.9°), C*α*-C-N (116.6° ± 1.3°), and in agreement to this work, the largest variability is observed for the bond angle N-C*α*-C. The bond angle values were compared to the B factor values averaged on the corresponding residues (Figure 1b). The lack of correlation between the values of bond angles and B factors shows that the variations of covalent geometry cannot be assigned to differences in protein internal mobility.

**Figure 1:**
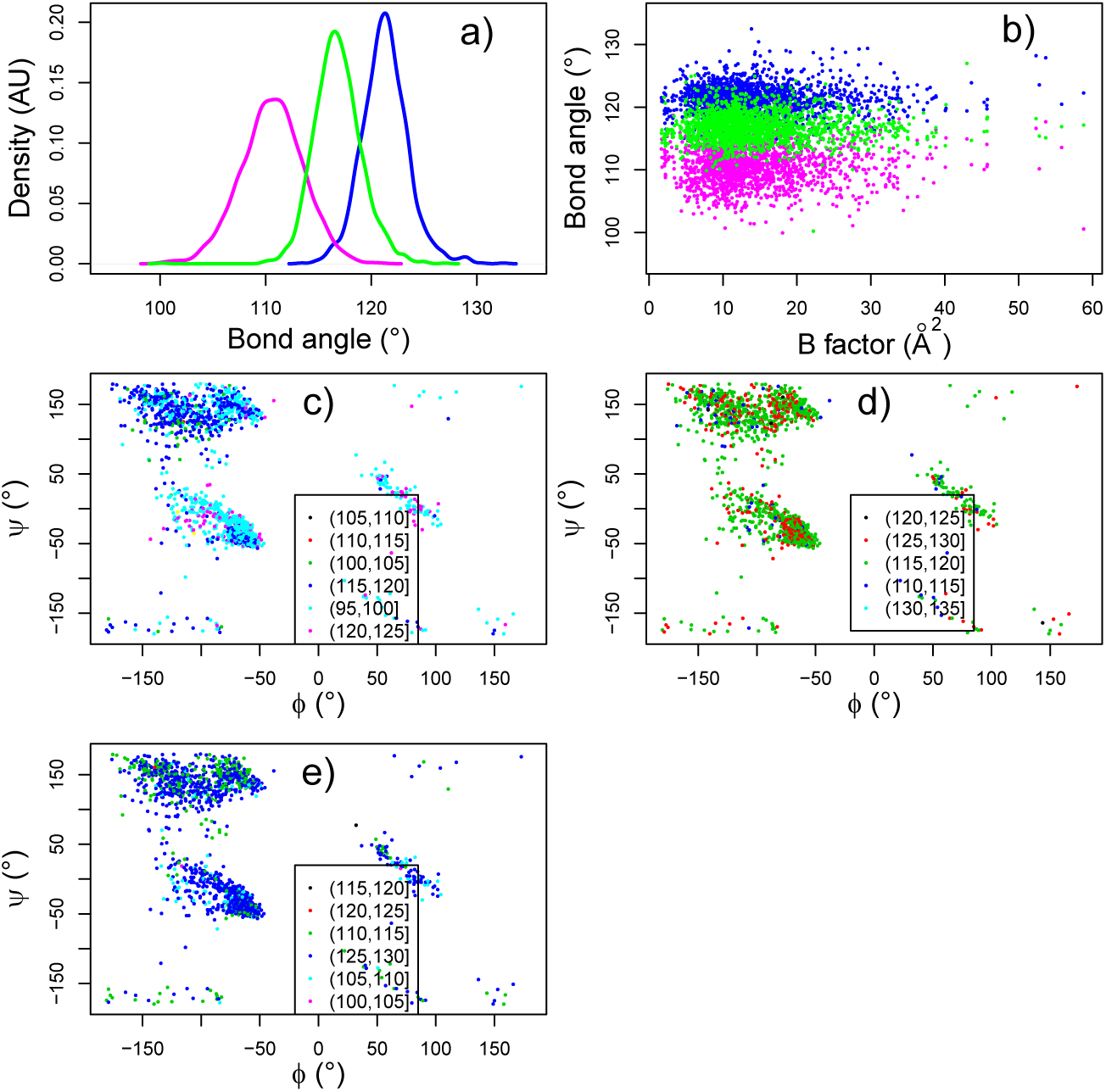
Variability of the covalent geometry within the 24 protein structures analyzed from the database top100.^61^ (a) Distribution of bond angles (°) between atoms C-N-C*α* (blue), N-C*α*-C (magenta) and C*α*-C-N (green). (b) Comparison between bond angles and B factors averaged on each residue. The color code is the same in (a) and (b). (c-e) Ramachandran plots of the protein structures colored according to the value of bond angles (°) between atoms C-N-C*α* (c), N-C*α*-C (d) and C*α*-C-N (e). On each plot (c-e), the color scale is defined in the inset legend.

The variations in covalent geometry were then plotted (Figure 1c-e) along the positions of protein residues in the Ramachandran diagram, by coloring the point describing the (*ϕ, ψ*) angle values of a given residue, according to the values of the residue bond angles. The Ramachandran plots are multi-colored according to the values of bond angles C-N-C*α* (Figure 1c), N-C*α*-C (Figure 1d) and C*α*-C-N (Figure 1e). All *α*-helix regions, around (−60°,-45°), display a quite monochromatic pattern, with values mostly in the range 100°-105° for angle C-N-C*α* (Figure 1c), in the range 125°-130° for angle N-C*α*-C (Figure 1d) and in the range 120°-125° for angle C*α*-C-N (Figure 1e). On the contrary, the *β*-strand region and the loops region of each diagram display a larger heterogeneity in bond angle values than the *α*-helix region. This heterogeneity has certainly a strong influence on the overall tertiary structures. Indeed, the *β* strands are extended structures in which local variations can have strong influence on the orientation at long distance. Similarly, the change of direction of protein backbone can be also very sensitive to local loop structure variation. In that way, both *β* strand orientations and loop directions have a strong impact on the protein tertiary structure.

In order to investigate the relevance of the uniform geometry hypothesis for the iBP calculation, the *ϕ*_angl_ and *ψ*_angl_ values measured on the top100 conformations have been compared to the *ϕ*_dist_ and *ψ*_dist_ values obtained as described in the Materials and Methods subsection “Input values for the calculation”. For each residue *K*, the cumulative sums of the differences between *angl* and *dist* backbone angles for residues *i, i* varying from 1 to *K*, were calculated:

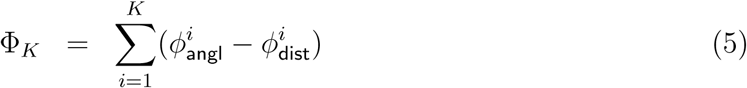

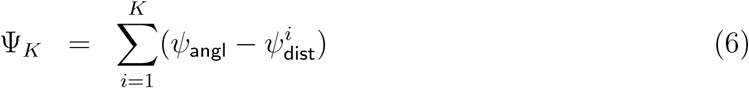

In Figure 2, the variations of Φ_*K*_ (green curves) and of Ψ_*K*_ (magenta curves) have been plotted along *K*, for the 24 studied proteins. The most important observation from these curves is that Φ_*K*_ and Ψ_*K*_ display extraordinary large variations along protein primary sequence. These variations extend from about 100° for the proteins 1benABH, 2bopAH, 1okAaH, bio1rpoH, up to several hundreds of degrees. The drift of Φ_*K*_ and Ψ_*K*_ depends of course on the total number of residues in the protein. Another observation is that Φ_*K*_ and Ψ_*K*_ curves do not display the same features. Φ_*K*_ curves are positive and increase along *K*, whereas Ψ_*K*_ curves are mostly negative and decrease along *K*. In addition, for most of the proteins, the comparison of the absolute values of Φ_*K*_ and Ψ_*k*_ reveals that one absolute value is larger than the other one, which induces a partial compensation between Ψ_*k*_ and Φ_*K*_ drift.

**Figure 2:**
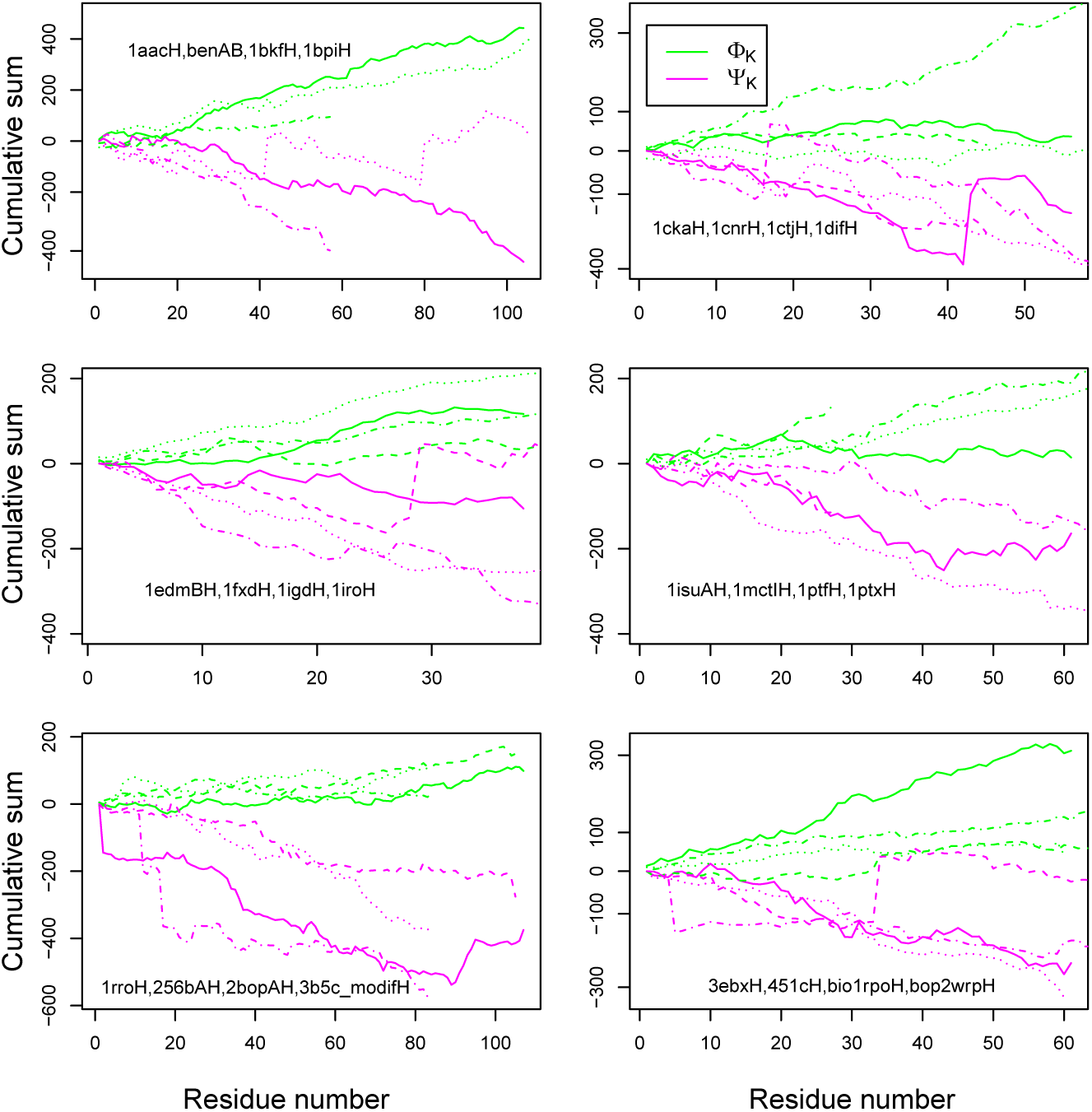
Variations of the Φ_*K*_ (green curves) and Ψ_*K*_ (magenta curves) (°) parameters (Eqs. 5 and 6) plotted along the primary sequences of target proteins. The entries of the analyzed proteins in the database top100^61^ are given on each plot.

To summarize, the analysis of protein structures involved in the present validation reveals that the hypothesis of uniform covalent geometry is far from being verified even in high-resolution crystallographic structures as the ones selected from the database top100^61^ with resolutions in the range 1.0-1.5 Å. Consequently, the differences between angles *ϕ*_angl_, *ϕ*_dist_ and *ψ*_angl_, *ψ*_dist_ display large cumulative drifts along the protein sequence.

### Exploring the conformational space of fragments using iBP

iBP calculations were performed on individual peptides spanning the analyzed proteins and the obtained results are presented in Figure 3. The run durations, plotted with decimal logarithmic scale, are centered around 10^2^-10^3^s, for error intervals of 20° (blue curve) and 40° (magenta curve), and jump to the 10^3^-10^4^s range for an error of 60° (green curve). The maximum run duration thus corresponds to about one day, which is not prohibitive.

**Figure 3:**
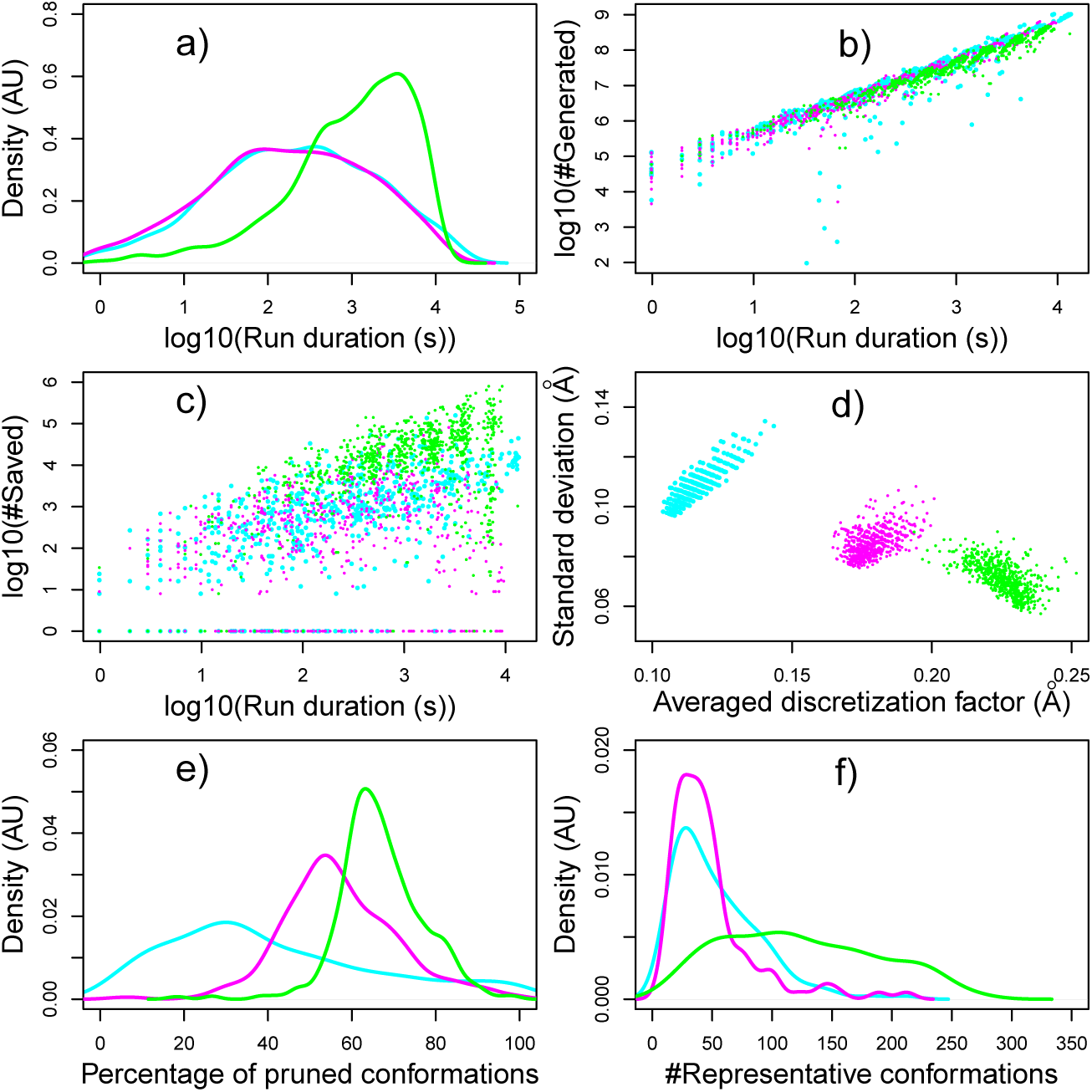
Analysis of the iBP runs on peptide fragments. The colors of lines/points correspond to the error interval on backbone angles: 20° (blue), 40° (magenta) and 60° (green). (a) Distribution of the run duration (s) plotted on logarithmic scale. Number of generated (b) and saved (c) conformations plotted along the run duration (s). Both axes are in decimal logarithmic scale. (d) Standard deviation (Å) of discretization factor along the average discretization factor (Å). (e) Distribution of the percentage of pruned conformations. (f) Distribution of the number of representative conformations obtained by clustering through self-organizing maps.

The tree sizes were reduced using the signed values of improper angles (Table 1), and are in the range 10^5^-10^9^. For each run, a maximal number of 10^9^ conformations to generate was required as input. The number of conformations saved during each run (Figure 3c) is in the range of 1 to 10^6^ which is several order of values smaller than this maximal number. All trees have thus been completely parsed during the iBP calculations.

The number of generated (Figure 3b) and saved (Figure 3c) conformations increase linearly along the run duration. The number of generated conformations is in the range 10^4^-10^8^, whereas the number of saved conformations is in the range 1-10^5^. For the largest error intervals (40° and 60°, magenta and green dots), similar numbers of conformations are generated (Figure 3b) and these numbers depend mainly on the duration of the run. In the case of the smallest interval width (20°, blue dots in Figure 3b), few runs display much smaller numbers of generated conformations. The number of saved conformations (Figure 3c) is of course smaller by two or three orders of magnitude from the number of generated conformations, but is also more dispersed. These numbers sample superimposed ranges for error intervals of 20° and 40° (blue and magenta dots), whereas they sample larger values for error interval of 60° (green dots). The result of tree parsing thus depends only slightly on the interval width, but the number of saved conformations vary qualitatively for error interval larger than 40° (Figure 3c, green points), and this difference is also visible in the run duration (Figure 3a, green curve).

The loss of information due to interval discretization during iBP calculations was analyzed (Figure 3d) through the discretization factor, which is the ratio between each distance interval to the number of tree branches generated within the interval. The standard deviation of this factor is plotted along its average value, both being calculated for the same individual iBP tree. Overall, one should notice that the largest discretization factors are smaller than Å. According to a recent work,^79^ the ensemble-average pairwise backbone RMSD for the microscopic ensemble underlying a typical protein X-ray structure is about 1 Å. The discretization factor of 0.25 Å is thus in the range of uncertainty of typical X-ray structures, and does not induce major loss of information in the iBP calculations.

For the various error intervals, the couples of average and standard deviation values for the discretization factor (Figure 3d) are clustered around different points: (0.11, 0.11) Å for the intervals of 20°, (0.18, 0.08) Å for the intervals of 40°, (0.23, 0.07) Å for the intervals of 60°. As expected, the average value increases with the interval width on *ϕ* and *ψ* angles. More surprisingly, the standard deviations decrease with the interval width: this is due to the discretization inputs. Indeed, the maximum number of branches is limited by 4 in all calculations, but the discretization factor should be always larger than a threshold of 0.05 Å. These two parameters induce the saturation of the number of branches for large widths as 60°. At the contrary, for smaller interval widths, the maximum number of branches is not attained for all distance intervals due to the required threshold. This induces a larger variability between the number of tree branches as well as a larger standard deviation.

During each iBP calculation, the conformations generated by branching on the *ϕ* and *ψ* intervals are then pruned or not according to the violation or to the verification of the distance interval between the C*α* atoms located at the N and C terminal residues. Percentages of pruned conformations (Figure 3e) are observed up to 100%. In the case of interval widths of 20° and 40°, three and one runs do not provide any solutions. As all these runs were performed using as input *ϕ*_dist_ and *ψ*_dist_ backbone angles, the pruning of all solutions is due to the inconsistency between the *ϕ* and *ψ* angle restraints and the extremities distance restraint. This inconsistency arises directly from the non-uniform covalent geometry described in the first section and is amplified by the use of a small error on backbone angle restraints.

Similarly to the number of runs without solutions, contrasted distributions are observed (Figure 3e) for the percentages of pruned conformations, depending on the width of intervals on backbone angles. For the smallest width (20°: blue curve), the percentage of pruned conformations displays a weak maximum at around 25%, but a non-negligible number of runs display percentages of pruned conformations in the 50-90%. Such high pruning percentages arise because in the case of narrow intervals on backbone angles, the hypothesis of uniform covalent geometry made by iBP has much more chances to induce solutions which do not verify the distance restraint between peptide extremities. For the larger interval widths on backbone angles (40°: magenta curve, 60°: green curve), the distribution is much more focused on larger percentages with respective ranges of 40-70% and 60-80%. The percentages larger than 80% are nevertheless vanishing for the largest interval widths. The global picture is that the increase of intervals on backbone restraints induces more pruning, but the percentage peaks at 50 and 60% obtained for interval widths of 40° and 60° are promising for the application of iBP to cases with error on restraints at the level of experimental cases. After generating peptide conformations using iBP, a procedure based on the self-organizing map^75^ is used to cluster the conformations and to extract representative ones. The distributions of the number of representative conformations (Figure 3f) are centered on the 0-100 and 0-50 range for the widths of 20° and 40°. Unsurprisingly, in the case of the larger width 60°, much larger numbers of representative conformations can be obtained, up to 250. The average number of representative conformations extracted from the SOM clustering of an iBP run on peptide fragment, is of the order of 10^2^, which makes the number of maximum combinations of peptides during the step of fragments assembly to be about 10^4^, and permits to overcome the combinatorial explosion, as it will be shown in the following.

### Efficiency of the TAiBP assembly strategy

Starting from iBP results, the individual peptide conformations were superimposed on the backbone atoms of their last and initial five residues, in order to grow the protein structure incrementally from the N terminal to the C terminal extremity. The proposed fragment assembly is then conserved or pruned according to two successively applied criteria: (i) the clashing criterion tests whether C*α* atoms of each fragment are farther apart from a given threshold (1 Å), (ii) the pruning distance criterion tests whether distance between the central C*α* of all inserted peptides is within 5Å of the distances observed in the initial PDB structure. Several assembly strategies have been used: (a) the fragments are added one by one from the N terminal to the C terminal extremities of the protein, (b) all possible assemblies of two fragments are formed along the sequence, and then assembled together successively from N to C terminal, (c) all possible assemblies of three fragments are formed along the sequence, and then assembled together successively from N to C terminal. Depending on the protein target, one approach can be more efficient than the others, but no general trend of efficiency for one strategy was found during the analysis, so the results of the three strategies are presented together.

Some statistics on the assembly steps of TAiBP are presented in Figure 4. The numbers, plotted in decimal logarithmic scale, of distance pruning events (#DistPruning), of clash pruning events (#ClashPruning) and of processed conformations (#Processed) are plotted along the number of peptide residues (Figures 4a-c). The numbers of distances (Figure 4a) and clash pruning (Figure 4b) events are mostly in the range 10^2^-10^4^ for all fragment sizes and interval of backbone restraints. In addition, these numbers increase of more than one order of value when the fragment size changes from 25 residues (assembly of two initial iBP fragments) to larger values. Larger pruning experienced in the case of larger fragments is probably induced by the excluded volume effect arising from the construction of the protein fold.

**Figure 4:**
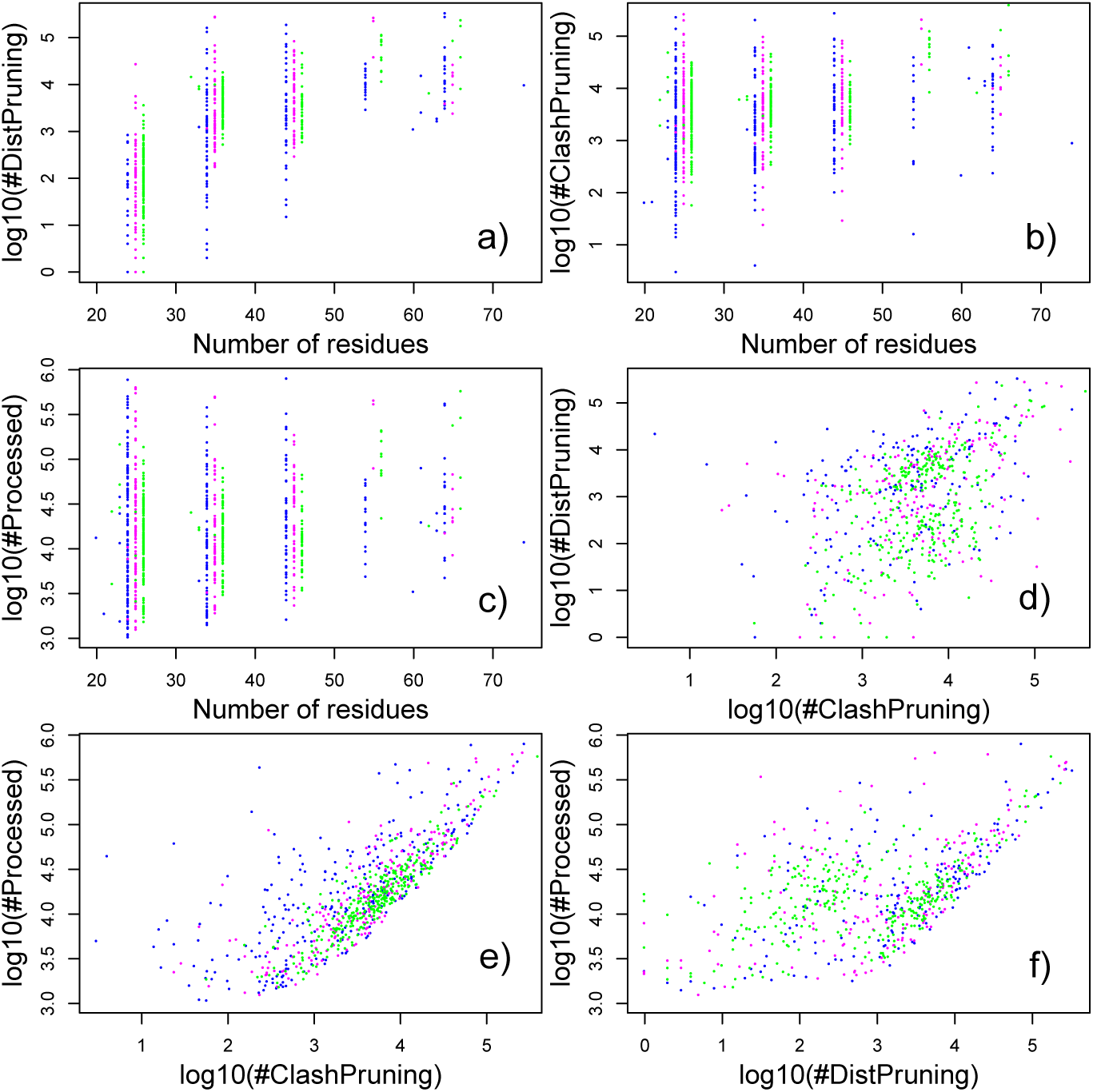
Analysis of the assembly of protein fragments to build the protein folds. All parameters except the number of residues in the assembled fragment are expressed in decimal logarithmic scale. The colors of points correspond to the error interval on backbone angles: 20° (blue), 40° (magenta) and 60° (green). The following plots are displayed: (a) number of distance pruning versus number of residues, (b) number of clash pruning versus number of residues, (c) number of processed conformations versus number of residues, (d) number of distance pruning versus number of clash pruning, (e) number of processed conformations versus number of clash pruning, (f) number of processed conformations versus number of distance pruning.

For fragment sizes larger than 25 residues, the numbers of pruning events (#DistPruning and #ClashPruning) are mostly around 10^3^-10^5^ (Figures 4a,b), in a range similar to the number of processed (#Processed) conformations (Figure 4c) which proves a large efficiency of pruning events for reducing the ensemble of solutions. Interestingly, distance pruning (Figure 4a) and clash pruning (Figure 4b) events are in similar range, displaying thus similar efficiency to reduce the number of solutions.

In the case of widths of 40 and 60° (magenta and green curves), the shifts observed for the parameters #DistPruning and #ClashPruning when increasing the interval widths on backbone angles, are steeper for the distance pruning events (Figure 4a) than for the clash pruning events (Figure 4b), whereas similar shifts are obtained for the interval width of 20° (blue dots). The widening of intervals has thus a stronger effect on the distance restraints between the peptide fragments than on the clash level. Finally, for fragments larger than 50 residues, the assembled fragments vanish except for the smaller intervals of 20° (blue dots), due to a pruning of all solutions. This pruning is the consequence of the discrepancy between non-uniform covalent geometry observed in the PDB structures and of the hypothesis of uniform covalent geometry made in the frame of iBP calculations.

The two by two comparisons of the events of distance and clash pruning, and of the number of processed conformations reveal the following trends (Figure 4d-f). The numbers of pruning events by clashes or by distances do not display any correlation (Figure 4d). At the contrary, #ClashPruning displays a quite strong correlation with #Processed (Figure 4e), specially for the largest angle interval (60°: green points). A similar tendency is observed for #DistPruning with two superimposed behaviors (Figure 4f): a correlation similar to the one observed for #ClashPruning, and other points with relatively smaller numbers of distance pruning events. This second set of points corresponds mostly to the case of the fragments of 25 residues. Indeed, as these fragments are much smaller than the full protein, they have less chance to be rejected by pruning distance information.

Each assembled fragment has been compared to the corresponding region in the top100^61^ target structure. This comparison was performed using RMSD (Å) between coordinates of heavy backbone atoms (Figure 5a,b) and not the TM score.^80,81^ Indeed, the statistical validation of TM score was performed on protein structures larger than 80 residues, ^81^ which do not correspond to the set of proteins studied here. For each fragment, maximum and minimum RMSD values are plotted in Figure 5a,b with respect to the fragment size.

**Figure 5:**
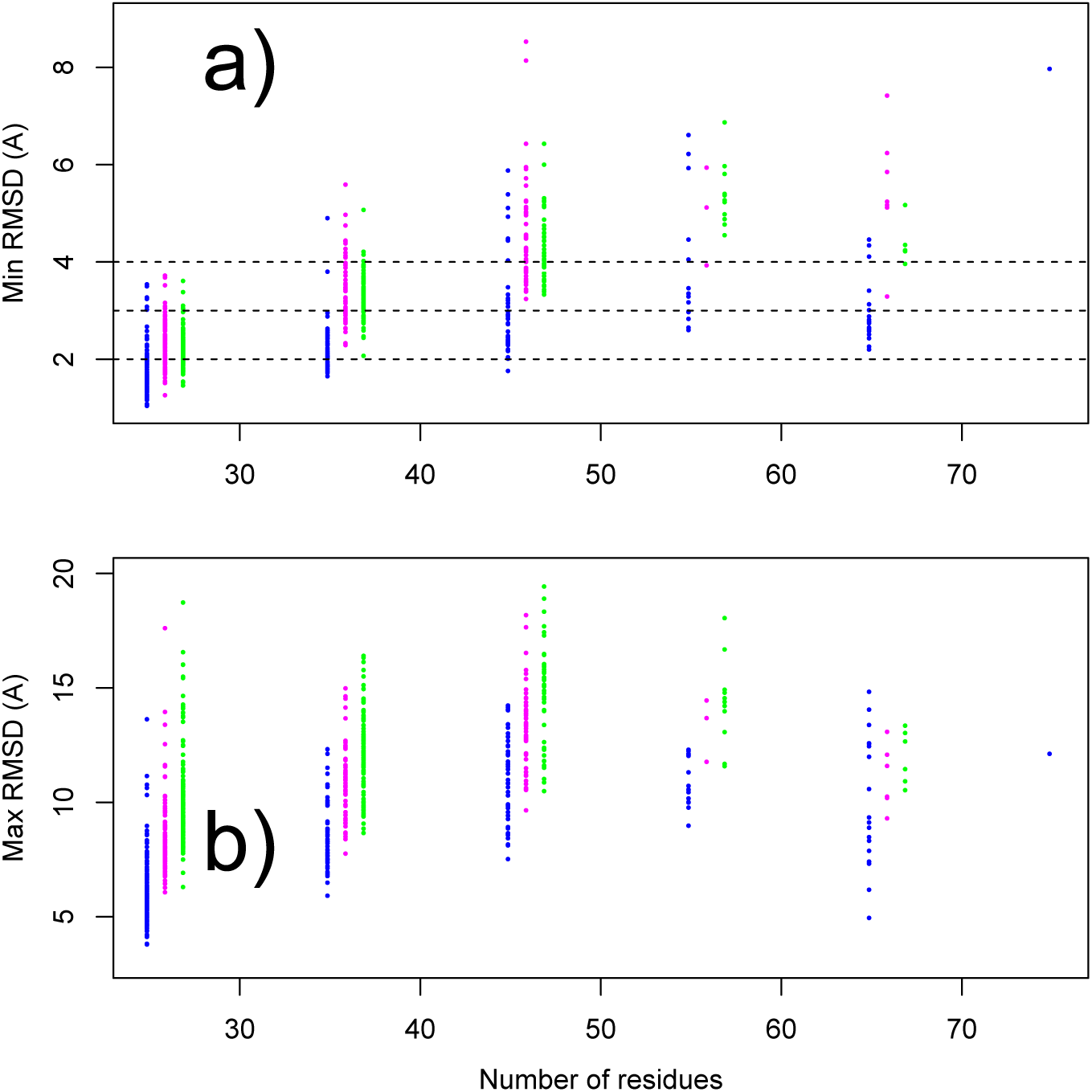
Comparison of the assembled fragment conformations to the corresponding regions in the top100 target structure. For each processed fragment, the minimum (a) and maximum (b) values of the coordinates RMSD (Å) between heavy backbone atoms are plotted along the number of residues. Dashed lines have been added at the levels of 2, 3 and 4 Å in minimum RMSD plot.

For each assembly calculation, the RMSD values to the target conformation were calculated for all TAiBP conformations. The maximum and minimum values of the RMSD distribution were then analyzed. In the case of narrow intervals (20°: blue points) on backbone angle restraints, minimum RMSD values are mostly smaller than 3.0 Å for all fragments up to 65 residues (Figure 5a). The increase of width in backbone angle intervals induces a drift of RMSD toward larger values: the RMSD drift is limited to 2-4 Å up to 35 residues, but jumps up to 5-6 Å for larger fragments. The threading-augmented iBP procedure proposed here thus allows one to obtain fragment conformations close to the PDB conformations for fragment sizes smaller than 65 residues. In that case, the Φ_*K*_ and Ψ_*K*_ drifts previously described (Eqs. 5-6 and Figure 2) have thus been overcome.

The maximum RMSD values (Figure 5b) are located in the 5-20 Å range. These maximum values were put in perspective with a previous analysis^82^ in which protein structures were compared to a representative set of protein-like alternative structures generated using threading. Most of the RMSD *R* values for an N-residue protein fall in the interval: 3.333*N* ^1*/*3^ *-* 2.0 ≤ *R* ≤ 3.333*N* ^1*/*3^ + 2.0, producing distributions of values smaller than 20 Å. This upper limit of 20 Å is similar to the one observed in the present calculation, which means that the TAiBP approach was able to mostly span the possible range of RMSD values. At the end of TAiBP calculation, the poly-Ala sequence was replaced by the protein specific sequence and the residue sidechains have been added using the relax tool of the Rosetta suite.^54^ The relax protocol consists of five cycles with rotamer repacking and minimization with progressively higher repulsive contributions within each cycle. ^83^ The obtained conformations have been analyzed (Table 3) and compared (Figure 6) to the conformation of the protein present in the database top100. From the 24 top100 structures initially processed, the 29 calculations realized on the 7 proteins smaller than 50 residues (Figure 6) display conformations calculated with TAiBP close to top100 conformations. Indeed, 18 runs display RMSD to initial top100 conformation smaller than 3 Å and 23 runs display RMSD smaller than 3.7 Å. Negative Rosetta total scores calculated according to Alford et al. ^84^ were obtained for all calculations, except the calculations on 1mctIH with an interval width of 60°.

**Table 3:**
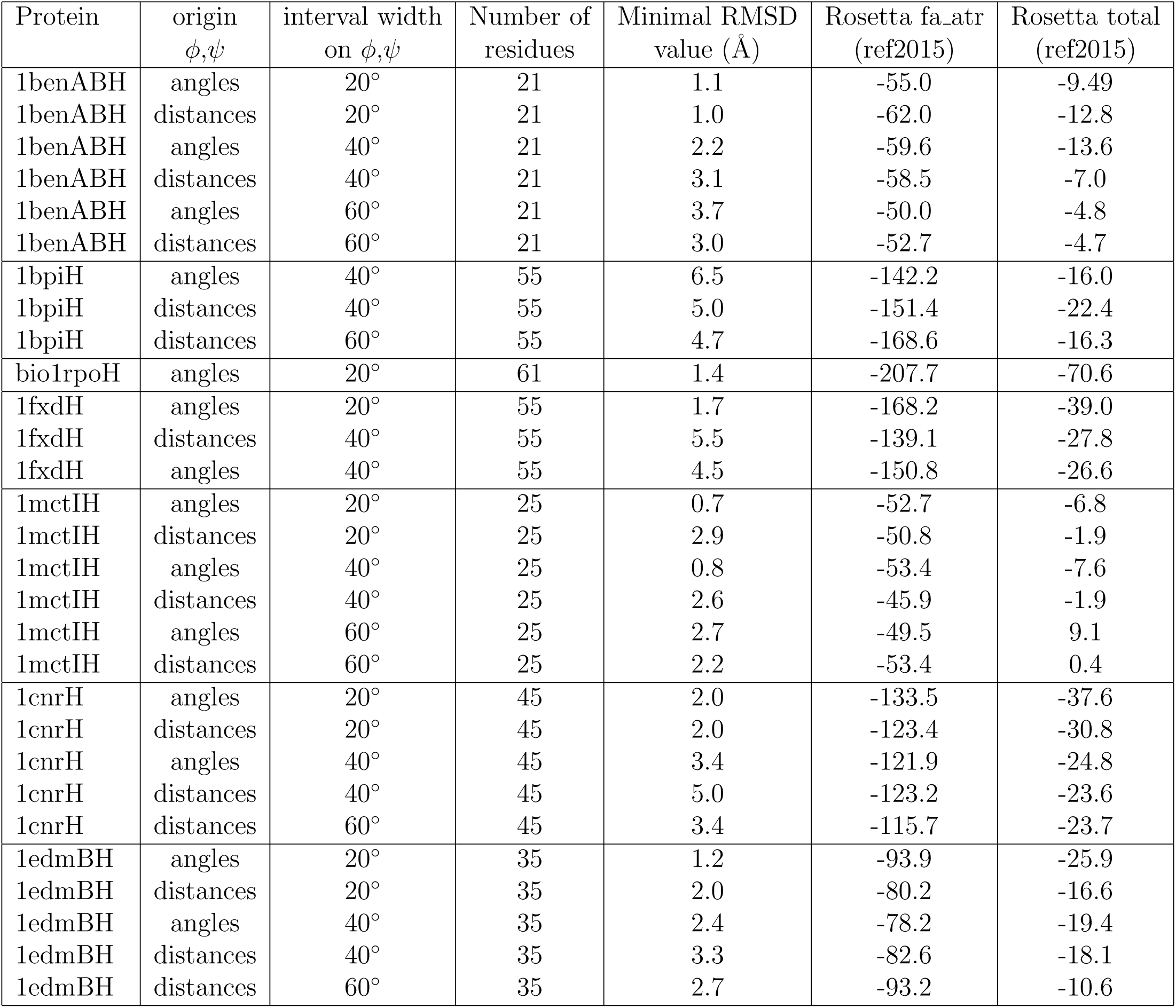
Analysis of the TAiBP conformations further relaxed using the Rosetta suite. ^54^ The Rosetta energy terms are defined from the ref2015 score function.^84^ Origin of *ϕ* and *ψ* angles describes whether their target values have been directly measured from the initial top100 conformation (angles) or whether they have been determined from distances measured on the top100 conformation using the hypothesis of a uniform covalent geometry (distances).

**Figure 6:**
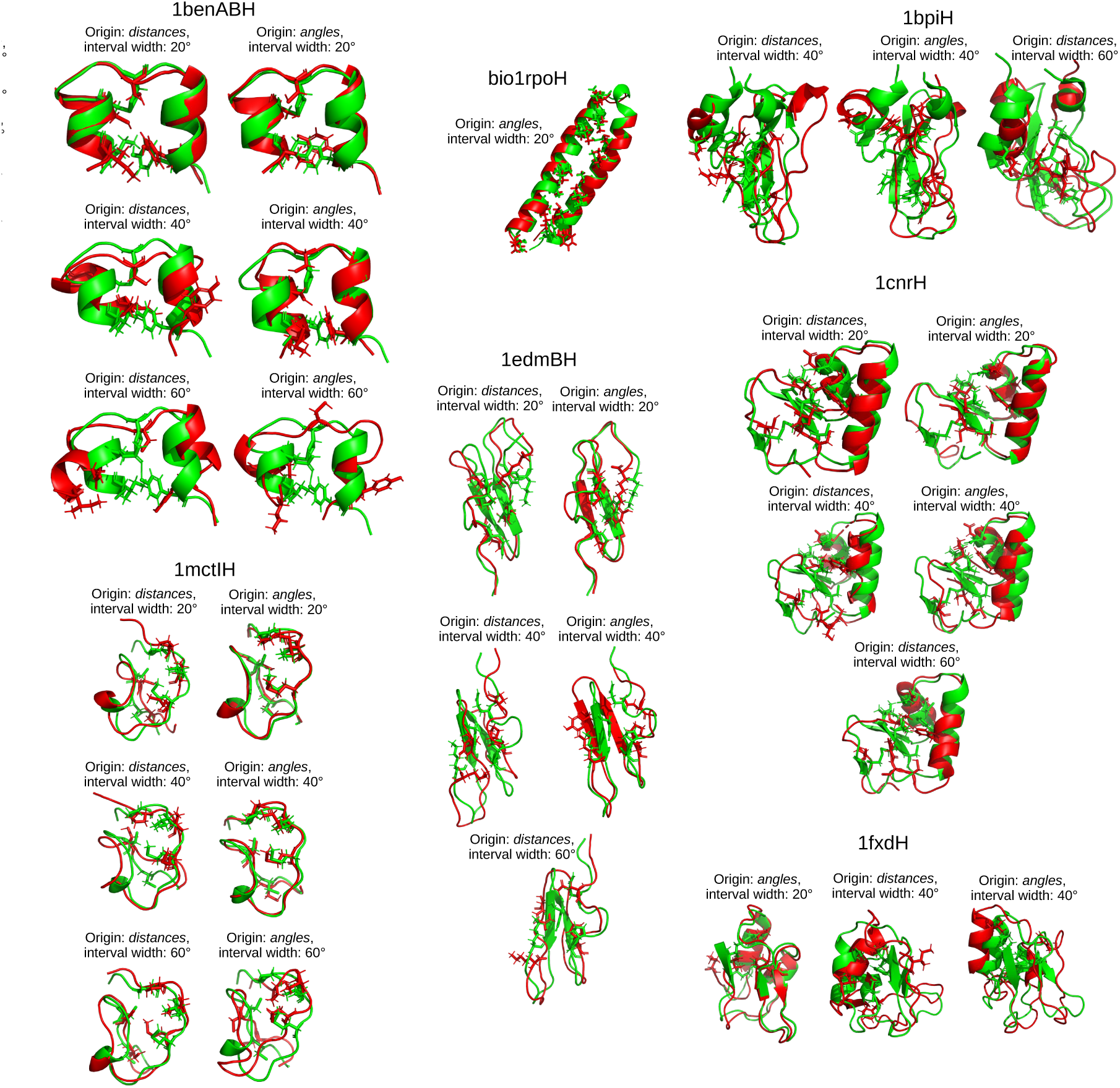
Comparison of the TAiBP conformations to the initial top100 conformations. The TAiBP conformations processed using the relax tool from Rosetta^54^ are drawn in red cartoon and superimposed to the target top100 conformations, drawn in green cartoon. Some residues of the protein core are drawn in licorice. Close to each superimposed structures, the origin of the backbone angles restraints *ϕ, ψ* is given as *distances* or *angles*. If the angles *ϕ, ψ* are measured on the initial top100 conformation, the restraints are of *angles* origin, whereas if the angles are calculated from measured distances in top100 conformations, assuming a uniform covalent geometry, the restraints are of *distances* origin. The interval widths of *ϕ, ψ* restraints are also given.

For a given protein, the origin of *ϕ* and *ψ* restraints: angles or distances, introduced in the subsection “Probing the hypothesis of uniform covalent geometry” display various influences (Table 3) on the coordinate RMSD between the TAiBP and top100 conformations. For 1ben1BH and 1bpiH, the RMSD is mostly smaller if the *ϕ* and *ψ* target values were extracted from the distances d(C^*i*^,C^*i*+1^) and d(N^*i*^,N^*i*+1^), assuming uniform covalent geometry. The entries 1fxdH, 1mctIH, 1cnrH and 1edmBH display an opposite trend.

The distribution of RMSD values, calculated on the whole sets of conformations obtained for a given TAiBP calculation (Figure 7) span values up to 10 Å. Due to the pruning events during the latest step of fragment assembly, this upper bound is smaller than the one observed for the maximum RMSD in Figure 5b. For the angles *ϕ*_angl_, *ψ*_angl_ measured on the initial top100 conformation, the increase of interval width induces a drift of RMSD toward larger values (magenta, brown and orange curves). The pattern is less clear for the angles *ϕ*_dist_ and *ψ*_dist_ calculated from measured distances in top100 conformations: in that case, many RMSD curves (blue, green and cyan curves) are more or less superimposed whatever is the interval width. The different proteins display quite different RMSD distributions which some distributions quite centered to a narrow interval and other much wider. These contrasted features arise from the various efficiencies of pruning long-range distances in the frame of different 3D protein topologies.

**Figure 7:**
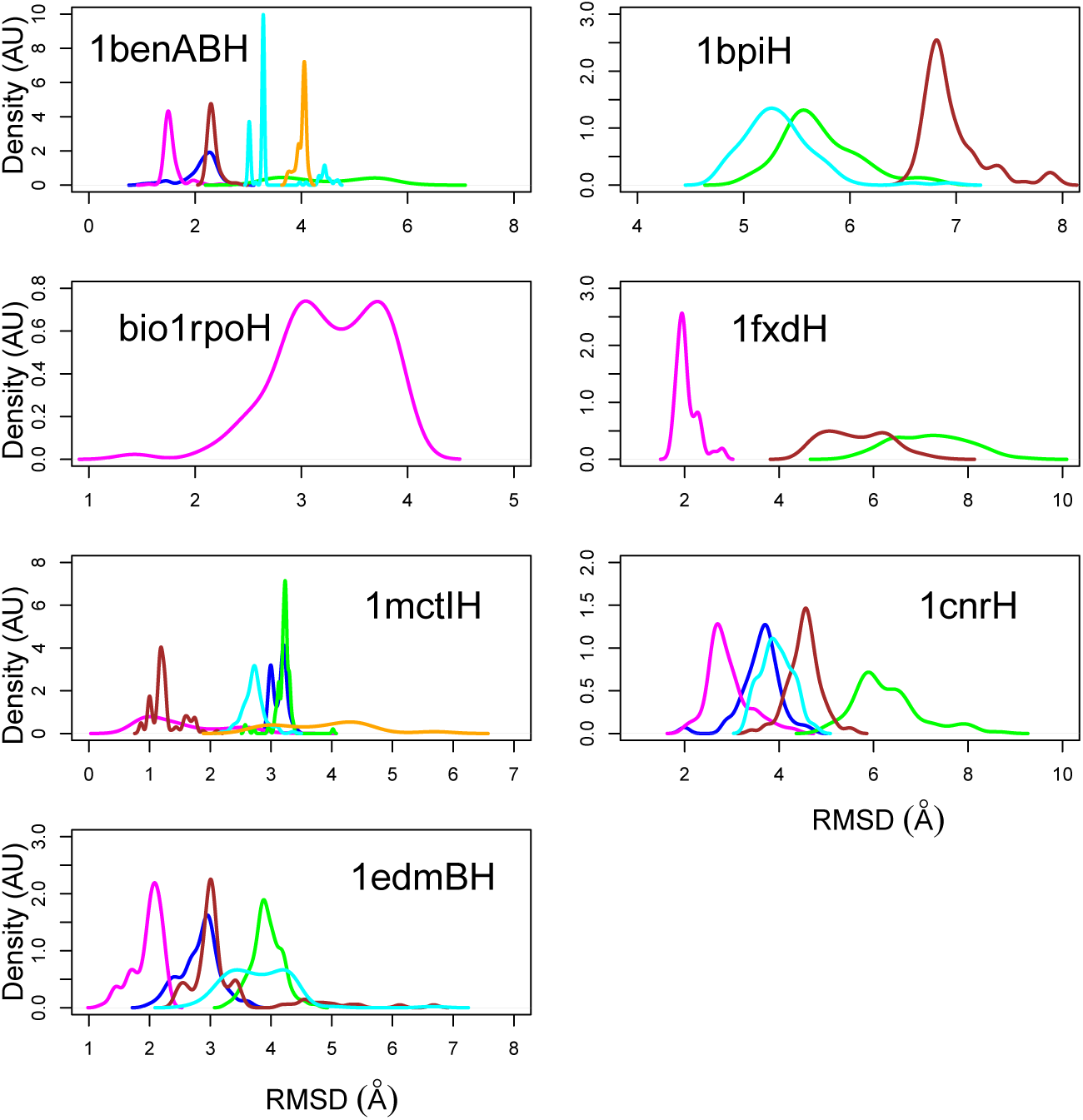
Distribution of RMSD values (Å) of each conformation obtained at the end of TaiBP approach with respect to the conformation of the corresponding protein in the database top100.^61^ The curves are colored according to the origin of the backbone angle restraints (*distances* or *angle*) introduced in the subsection “Input values for the calculation” in Materials and Methods, and to the interval of *ϕ, ψ* restraints. The color used are: blue (*distances*, 20°), magenta (*angles*, 20°), green (*distances*, 30°), brown (*angles*, 30°), cyan (*distances*, 40°), orange (*angles*, 40°).

### Influence of uniform covalent geometry

The covalent geometry of the 1benABH, 1cnrH, 1edmBH, 1fxdH, 1igdH, 1isuAH, 1mctlH, bio1rpoH and 1bpiH conformations obtained using TAiBP and then relaxed using Rosetta displays some characteristics (Figure 8) quite different with respect to the ones analyzed on the top100 conformations at the beginning of the present work (Figure 1). Similar trends in covalent geometry were observed in conformations obtained by TAiBP before adding sidechains and relaxing (data not shown). In the relaxed conformations, the distribution of covalent angles (Figure 8a) is much thinner, although some individual deviations are observed (Figure 8b). These large drifts are observed in the protein regions in which two neighboring peptide fragments were superimposed. The *ω* dihedral values (Figure 8c) are distributed around 180° and −180°, in a similar way than in the initial structures (data not shown). These variations of *ω* dihedral angles have been already observed from various analyses of X-ray crystallographic structures. ^65–67^

**Figure 8:**
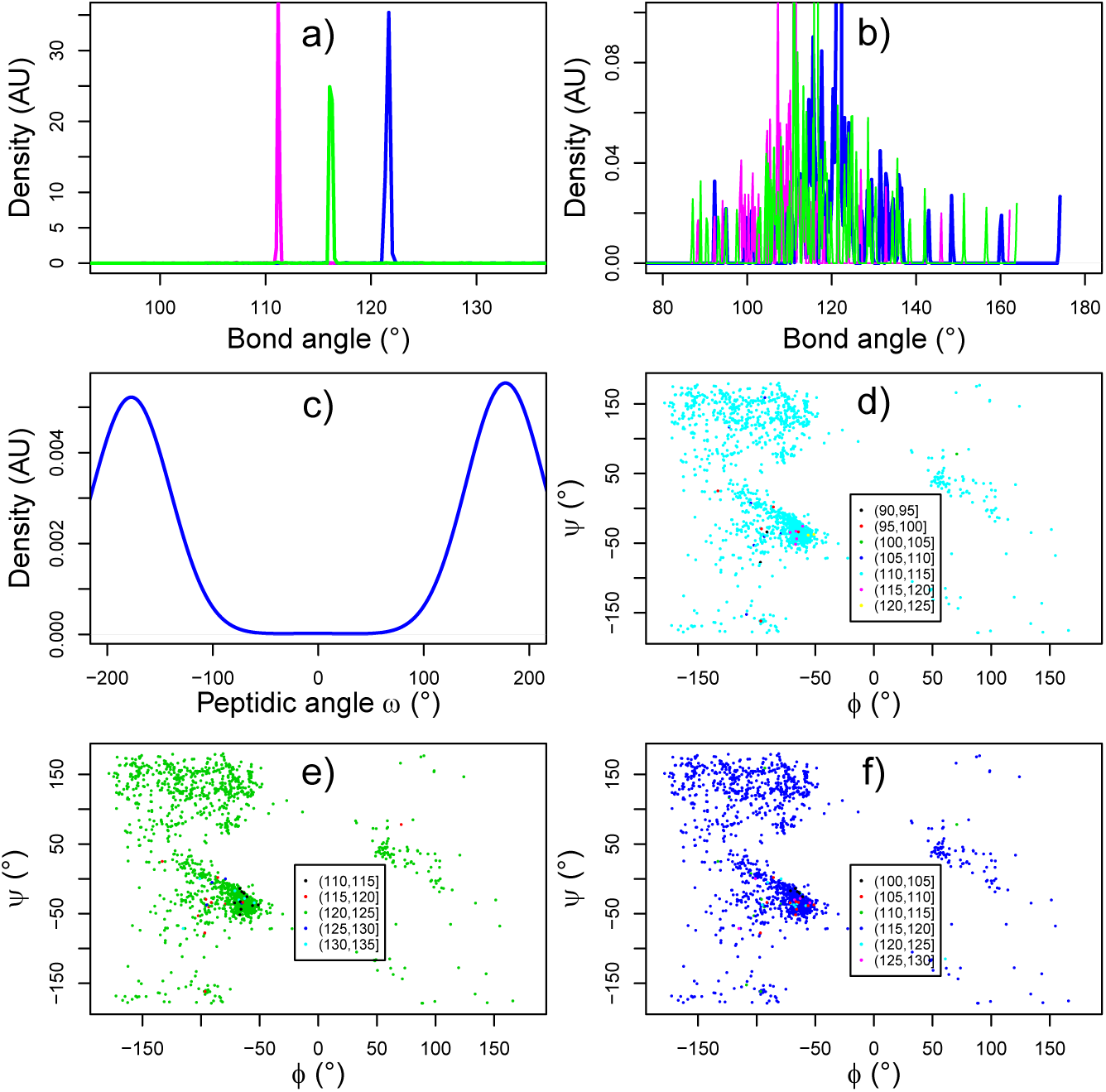
Variability of the covalent geometry within the set of the conformations obtained on 1benABH, 1cnrH, 1edmBH, 1fxdH, 1igdH, 1isuAH, 1mctlH, bio1rpoH, 1bpiH using TAiBP and then relaxed using Rosetta. (a) Distribution of bond angles (°) between atoms C-N-C*α* (blue), N-C*α*-C (magenta) and C*α*-C-N (green). (b) Distribution of bond angles (°) between atoms C-N-C*α* (blue), N-C*α*-C (magenta) and C*α*-C-N (green) with increased scale to detect the outlier peaks, (c) Distribution of the *ω* dihedral angle (°). (d-f) Ramachandran plots of the protein conformations colored according to the value of bond angles (°) between atoms C-N-C*α* (d), N-C*α*-C (e) and C*α*-C-N (f). On each plot, the color scale is defined in the inset legend.

The comparison of the Ramachandran plots between Figures 1 and 8 reveal two differences. First, the TAiBP Ramachandran plots display a narrower range of colors than the top100 ones in agreement with the thinner distribution of covalent angles. Second, the (*ϕ,ψ*) distributions are fuzzier in Figures 8d-f than in Figures 1c-e. The convergence toward a uniform covalent geometry is thus accompanied by a expansion of the regions sampled in the Ramachandran diagram. Such expansion of allowed regions has been also observed in a recent analysis of the Ramachandran diagram.^85^ In agreement with covalent geometry close to uniformity, the plots of cumulative sum of the differences: Φ_*K*_ (Eq. 5) and Ψ_*K*_ (Eq. 6) display much smaller drifts on the TAiBP conformations with a large majority of the values ranging between −50° and 50° (data not shown).

The protein conformations generated by the TAiBP approach and relaxed with Rosetta (Table 3 and Figure 6), have been used as target conformations for a new run of the TAiBP approach, in order to investigate whether, in the case of mostly uniform covalent geometry, different results could be obtained. In a way similar to the previous TAiBP run, the coordinate RMSD (Å) between the new target and TAiBP fragments display a drift toward larger values for increasing fragment size (data not shown). The minimal and maximal RMSD distributions calculated for the reconstructed full chains of protein targets (Figure 9) show that for all targets except 1fxdH, the distribution of minimal values are mostly in the 1-4 Å for the interval width of 20° (blue curves). For all targets except 1bpiH and on a lesser extend 1cnrH, the increase of interval width do not have strong impact on the minimal RMSD distribution (full lines). Using target conformations closer to the hypothesis of uniform covalent geometry thus reduces the impact of increased intervals for *ϕ* and *ψ* angles. The distributions of maximal RMSD values (dashed lines) displays more variability than minimum RMSD distributions. Unsurprisingly, these distributions shift toward larger values and/or become broader in the case of increased interval width for *ϕ* and *ψ* values.

**Figure 9:**
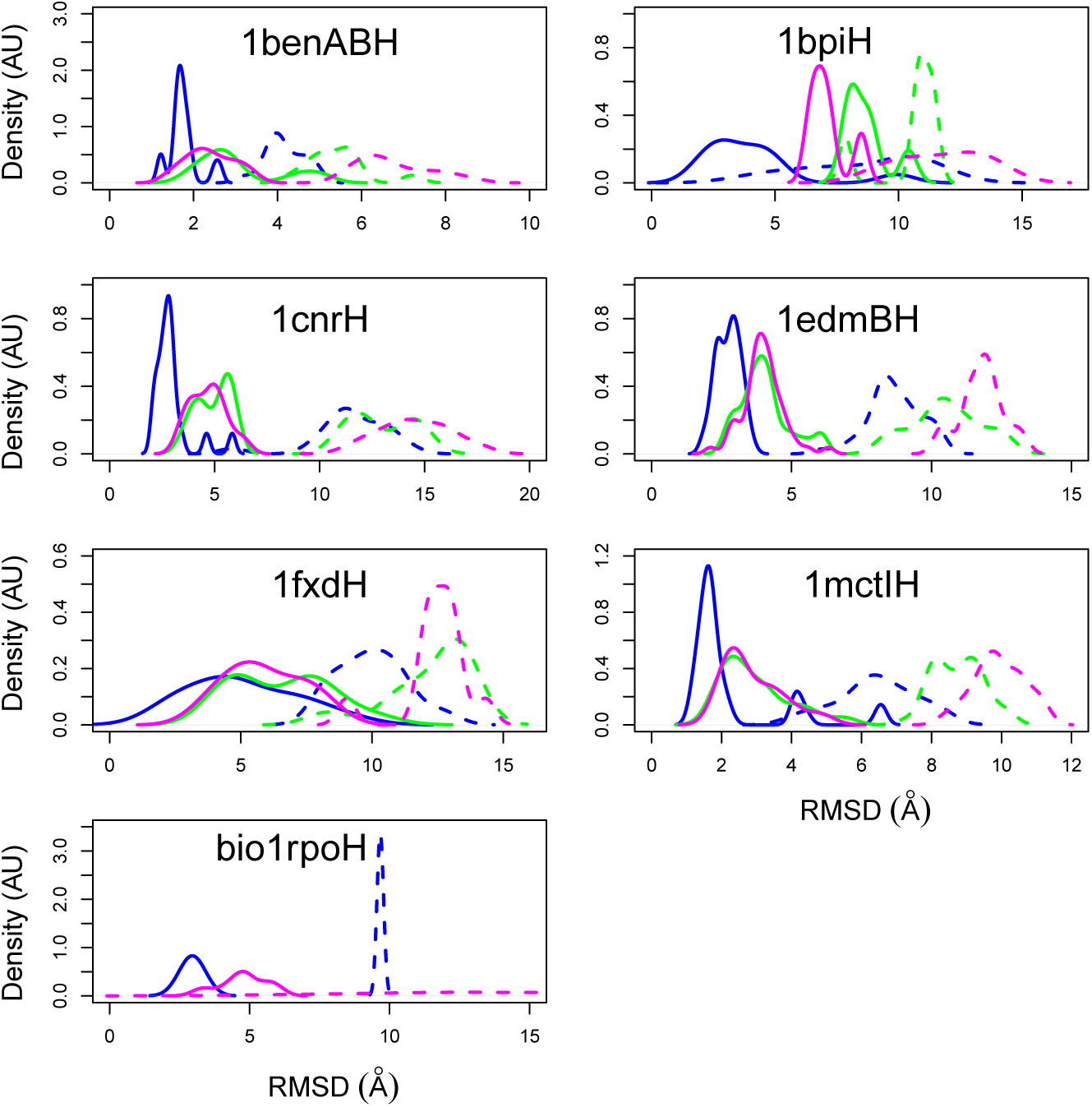
Distribution of minimum (full line) and maximum (dotted line) RMSD (Å) values between the target and the TAiBP protein conformations. These RMSD were obtained during a TAiBP run using as targets protein conformations generated by the TAiBP approach and relaxed with Rosetta (Table 3 and Figure 6). The calculations were realized with interval of 20° (blue), 40° (green), and 60° (magenta) widths on the *ϕ* and *ψ* values.

## Discussion-Conclusion

Two main conclusions can be derived from the present work. The most important one is the design of an approach allowing to systematically parse all conformations of a protein up to 100 residues, using low precision input restraints. The other outcome of the manuscript is that the variability of covalent local geometry is an essential parameter for building protein conformations. Until now, this aspect has been discussed only very little in the literature.^78^

A method has been described to generate the protein structure by systematically exploring all possible conformations of the protein. This method is based on a threading-augmented interval Branch-and-Prune (TAiBP) approach in which the interval Branch-and-Prune (iBP)^28,44,86^ is first used to systematically explore the conformations of 15-residues peptides fragments of the protein, followed by the construction of protein structure by systematically assembling fragment’s conformations, and by pruning conformations displaying atom steric clashes and violations of few long-range distance restraints.

This two steps approach along with clustering using self-organized maps^75^ allows to overcome the combinatorial explosion arising from the exponential complexity of the iBP algorithm. The duration of a total calculation is of the order of tenths of hours. This could be even speed up by using compiled language in place of python scripts used for fragments assembly and clustering.

The calculations performed using the TAiBP were validated by detecting whether this approach provides at least one solution close to the target solution. This detection was performed using the coordinate RMSD between backbone atoms. For most of the proteins smaller than 50 residues and interval widths of 20° for backbone angles, solutions were obtained with RMSD values smaller than 4 Å. Larger protein sizes and/or larger interval widths induce drift in the obtained conformations, which usually conduct to a pruning of all conformations because the long-range distance restraints are no more verified.

The largest problem to which the proposed approach faces is the non-uniformity of covalent geometry among the PDB structures. This aspect of PDB structures looks very minor as it involves only few degrees variations among the covalent angles, but, as this non-uniformity induces biases in the direction of extended and loop parts of protein structures, it is obvious that it may have big consequences in fragment assembly. Also, as this non-uniformity is present since the first days of structural biology and is probably deeply related to amino-acid type and to the position in the Ramachandran diagram,^66,87–92^ it is quite difficult to sort it out. Nevertheless, it should be noticed that attempts have been made^93^ to explore the relationship between backbone conformations and covalent geometry.

On the other hand, the iBP approach on which we based the conformational sampling of protein fragments, was developed using the initial hypothesis of a uniform covalent geometry. This hypothesis permitted to set up an algorithm which displays good scaling properties in case of a sufficient number of exactly known inter-atomic distances.^28^ Modifying the algorithm to take into account possible variations in the covalent geometry would increase enormously its complexity. Nevertheless, the very recent development of residue-specific force fields^37–40^ opens new avenues for taking into account these aspects.

The present work opens the way to the use of iBP approach for exploring systematically the conformational space of proteins, using geometric restraints analogous to those experimentally measured by NMR. The validation of this systematic exploration of protein conformational space deeply changes the perspectives on protein structure calculation.

## Acknowledgments

Thérèse Malliavin thanks the Pasteur Foundation for postdoctoral support of Dr Bradley Worley and thanks Dr Bradley Worley for its implementation of iBP and Dr Guillaume Bouvier for his support in python scripting. The authors wish to thank Institut Pasteur, CNRS, Ecole Polytechnique, FAPESP, CNPq and the program Infinity of CNRS for financial support.

